# Stainless Steel vs Calcium Fluoride for biomedical Raman Spectroscopy

**DOI:** 10.64898/2026.10.09.753418

**Authors:** Cynthia Ng, Nathan Blake, Geraint Thomas

**Affiliations:** Department of Cell and Developmental Biology, University College London, Gower Street„ London, WC1E 6BT, UK; Department of Medical Physics and Biomedical Engineering, University College London, Gower Street„ London, WC1E 6BT, UK; Department of Informatics, King’s College London, Bush House, Strand Campus, 30 Aldwych, London WC2B 4BG, UK

**Author notes:** These authors contributed equally to this work (Cynthia Ng, Nathan Blake).

## Abstract

Raman spectroscopy (RS) has been shown to differentiate various disease states in tissue samples, yet the impact of sample substrate on spectral quality and downstream machine-learning (ML) performance is under-explored. We collected 960 spectra from 36 formalin-fixed, paraffin-embedded colonic specimens mounted in parallel on calcium fluoride (*CaF*_2_) and mirrored stainless-steel (*SS*) slides. Raw *SS* spectra were more intense than *CaF*_2_, and routine dewaxing left prominent paraffin peaks on *CaF*_2_ that were largely absent on *SS*. Extending *CaF*_2_ xylene immersion to 48 h reduced key paraffin peaks by 11–23 %, aligning its biochemical profile with *SS*. Nested cross-validated classifiers were built to distinguish true disease (T1) from epithelial misplacement (EM). The highest accuracies were observed on *SS* (0.65–0.66), while *CaF*_2_ improved from 0.41–0.57 to 0.65–0.69 after extended dewaxing, chiefly via gains in recall. Although optimal dewaxing renders *CaF*_2_ diagnostically non-inferior, *SS* still offers superior accuracy, faster preparation, at approximately 4% of the cost, supporting its integration into scalable, AI-enabled RS pathology workflows.

## 1 Introduction

Raman spectroscopy (RS) has many biomedical applications, including the classification of tissues for diagnostic purposes ^1,2^. However, there remains several elements of the process which would benefit from optimisation^3^. One of these is the sample substrate: the physical surface on which a sample is mounted. The standard substrate for tissues is calcium fluoride (*CaF*_2_). This benefits from low autofluorescence, which can often overwhelm Raman signals and is not Raman active in the relevant wavenumber region ^4^. However, it is approximately three orders of magnitude more expensive compared to standard glass pathology slides, and is more brittle, resulting in occasional breakages and reducing the suitability for routine lab use. Stainless steel (*SS*) is much more robust and has been proposed as an alternative approximately at 4% the cost of *CaF*_2_. It has also demonstrated enhanced Raman signal intensity, improved signal-to-noise ratios (SNR), accelerated spectral acquisition times due to a double-pass photon reflection mechanism ^5^, and more complete paraffin removal^6^.

Despite this early promise of SS, *CaF*_2_ remains a common choice. In part, this may be because the observed benefits of *SS* have not been shown to translate into improved clinical outcomes. In this study, we compare the analysis of the same clinical data set mounted on both *CaF*_2_ and *SS*, evaluating whether either substrate demonstrates higher signal acquisition or improved paraffin removal. Further, for the first time in the literature, we assess whether any such improvements actually translate to superior machine learning (ML) performance, which could lead to improved clinical outcomes.

## 2 Experimental Set-up

### 2.1 Clinical problem

Epithelial misplacement (EM) is a phenomenon whereby dysplastic glands from a colorectal adenoma are displaced into the submucosa. This is often attributed to mechanical distortions such as polyp torsion, biopsy, or endoscopic manipulation, which disrupt the muscularis mucosae and allow glandular elements to herniate into deeper layers of the bowel wall^7^. This benign phenomenon can closely mimic submucosal invasion, complicating histological diagnosis, risking overtreatment or misclassification of malignancy.

RS may have the biochemical sensitivity required to distinguish EM from true malignancy. In this study we obtained 36 samples from 35 patients. 18 samples were EM and 18 were T1 colorectal cancer, confirmed by two specialist gastrointestinal (GI) pathologists.

### 2.2 Sample and Substrate Preparation

Formalin-fixed, paraffin-embedded (FFPE) colonic tissue samples were acquired from UCL/UCLH Biobank (Ethical Approval: REC 25/YH/0120). Adjacent sections were cut at 8 µm and mounted onto calcium fluoride (*CaF*_2_) and stainless steel (*SS*) slides. Additional adjacent sections were cut at 3 µm and mounted onto glass slides for standard H&E staining. Regions of interest (ROIs) were identified by an expert GI pathologist based on the H&E slides.

Dewaxing was undertaken in two stages. The initial protocol involved sequential immersion in several changes of xylene over a 40 minute period for both *SS* and *CaF*_2_ slides, followed by an extended immersion for 24 hours for the *CaF*_2_ slides only, after which all slides were rehydrated using graded ethanol steps^6^. Spectra were acquired from these samples as described in Section 2.3. A further dewaxing step was then applied to the *CaF*_2_ slides using an extended xylene immersion of 48 hours to optimise paraffin removal. The differences in dewaxing efficacy between substrates were subsequently evaluated through spectral analysis.

### 2.3 Raman set-up

Spectral data were acquired using the Renishaw inVia Qontor Raman microscope equipped with a 785*nm* laser and a 50*x* objective lens (Renishaw plc, Wotton-under-edge, UK). Point-scanning mode was used, with each measurement consisting of single-point acquisitions within the designated ROIs. Calibration was performed before each imaging session using a silicon standard to ensure spectral consistency, with the characteristic silicon peak at 520.5*cm*^*−*1^ as reference. Comparable tissue sections were Raman mapped on the same day with the same calibration readings.

An integration time of 10 seconds was used for all measurements, with each spectral acquisition comprising 10 accumulations per point. A total of 10 points were recorded per sample, yielding approximately 20 spectra per region. Overall, 960 spectra were collected, 320 for each of the three substrate conditions (*SS*, initial dewaxed *CaF*_2_ and extended dewaxed *CaF*_2_).

### 2.4 Analysis and machine learning

Unless stated otherwise all acquired spectra were processed using code written in Python 3.10. Preprocessing involved the removal of cosmic ray signals via manufacturer-provided algorithms, truncation to the fingerprint region (500 – 1800*cm*^*−*1^), and standard-normal variate normalisation (SNV). Mean spectral overlays were generated for both unnormalised and SNV-normalised data, with key Raman bands identified and compared across substrates.

Standard machine learning algorithms were employed to compare performance of discriminating between EM and T1 samples across the four substrate conditions. These were principal component analysis - linear discriminate analysis (PCA-LDA), support vector machine (SVM) with a radial basis function kernel and gradient boosting (GB).

#### 2.4.1 Nested Cross Validation

Each ML method has associated hyper-parameters and these benefit from optimisation. We performed a limited grid-search over the hyper-parameter space within a nested cross-validation (CV) framework. This involved 5-fold CV in the outer fold and 3-fold CV in the inner fold 1, the latter allowing hyper-parameter optimisation without over-fitting. All splits were performed at the level of the sample, and was stratified to ensure approximately equal labels. These strategies mitigate over-fitting, resulting in more realistic model performance estimates^1^.

#### 2.4.2 Paraffin Contamination Assessment

To evaluate residual paraffin contamination, spectra collected from pure paraffin samples on both *CaF*_2_ and *SS* slides were used as and authentic standard reference. Prominent paraffin-associated peaks at 1062, 1132, 1170, 1295, 1417, 1439, and 1462 cm^*−*1^ were used to identify and assess contamination levels across tissue spectra^6^.

Two datasets were initially analysed: (1) a preliminary dataset collected after the initial dewaxing step, where *CaF*_2_ spectra (labelled “CaF”) still showed significant paraffin contamination, and (2) an extended dewaxing dataset (“CaFII”), where spectra were recollected after an additional 48-hour xylene treatment to enhance paraffin removal.

Differences in peak intensity were also compared between *CaF*_2_ and *SS* slides. Notably, *SS* substrates appeared to facilitate more complete paraffin removal during standard dewaxing and may also induce paraffin melting under laser exposure, contributing to the observed spectral differences.

## 3 Results

### 3.1 Spectral acquisition and signal intensity comparisons

Raman spectra were acquired using identical experimental parameters for both *CaF*_2_ and *SS* substrates. To maximise signal strength on *CaF*_2_, laser power was increased to 100% with 10 accumulations per point, following optimisation of acquisition settings. Nevertheless, spectra collected from *CaF*_2_ displayed consistently lower signal intensity than those from *SS*. As shown in figure 2, the overlay of unnormalised mean spectra demonstrates that *SS* yielded consistently stronger peaks across the entire Raman shift range. In particular, higher intensities were observed at key wavenumbers corresponding to biomolecular features—such as amide I (1660 cm^*−*1^) and amide III (1250 cm^*−*1^) for proteins, 1445 cm^*−*1^ for lipids, and 1085 cm^*−*1^ for nucleic acids.

**Figure 1.**
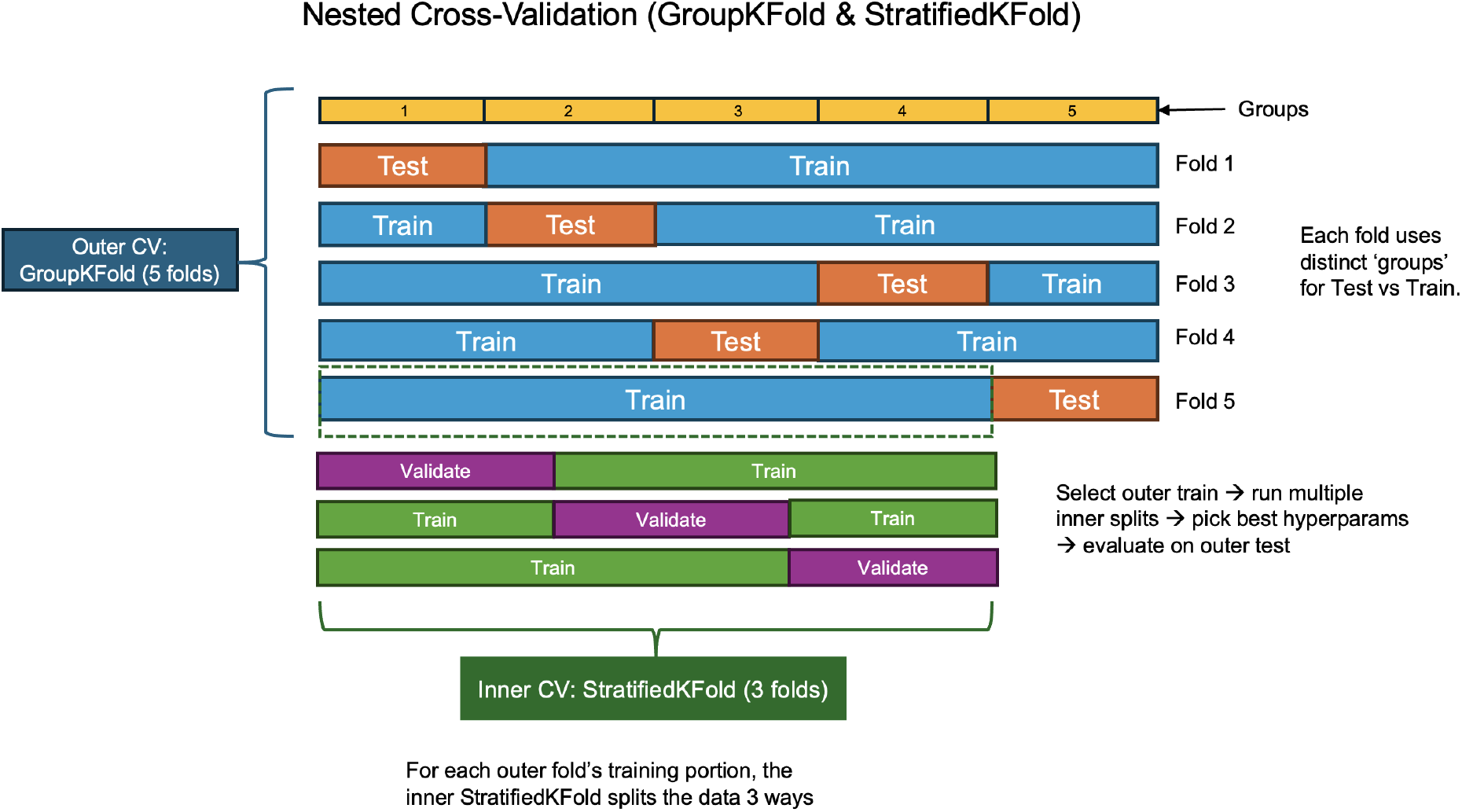
Nested cross-validation strategy with `GroupKFold` (outer) and `StratifiedKFold` (inner).

**Figure 2.**
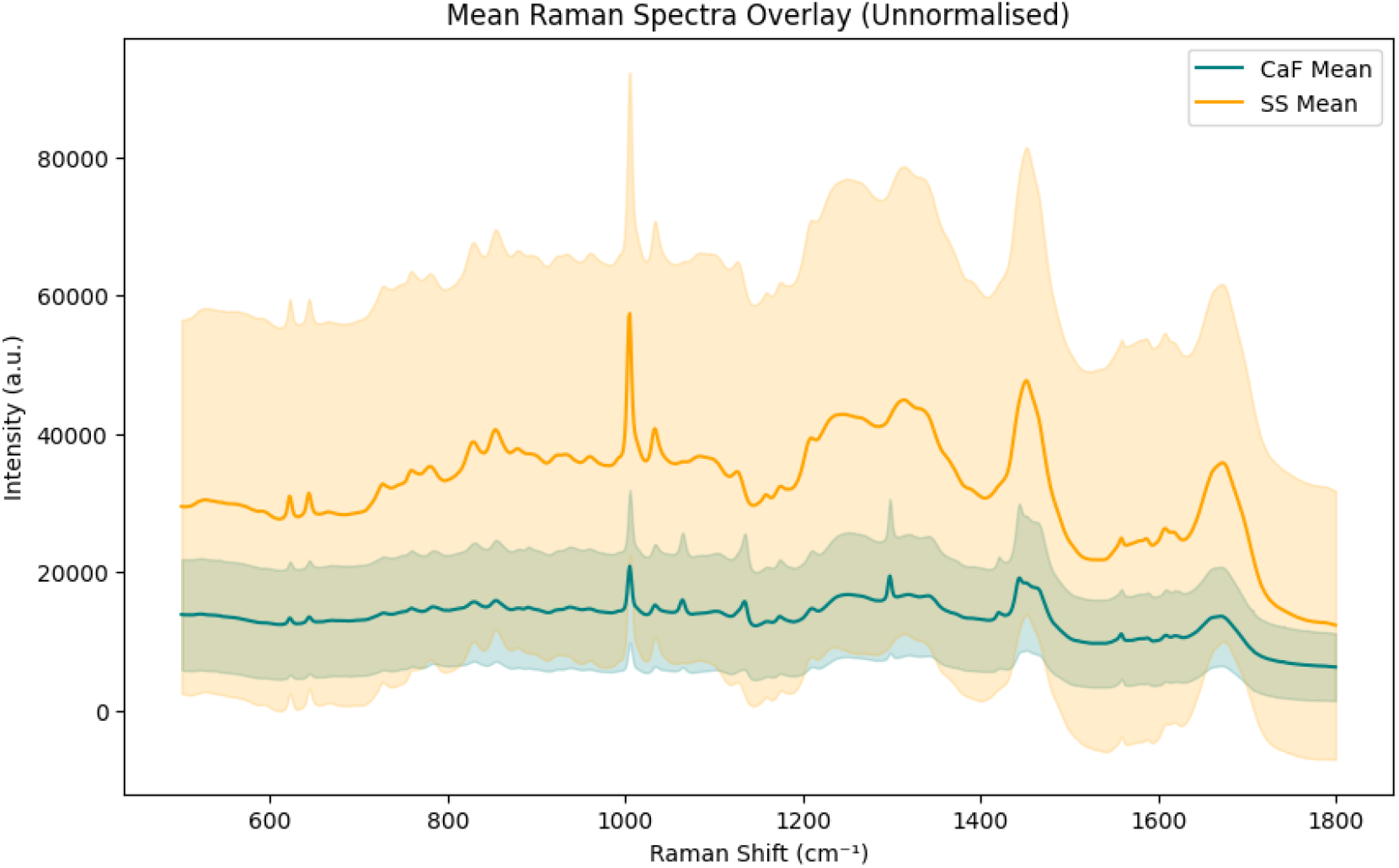
Mean unnormalised Raman spectra overlay from *CaF*_2_ and *SS* substrates, with standard deviation

To determine whether the observed differences were due solely to intensity variation or also reflected biochemical disparities, SNV normalisation was applied to the spectra. After normalisation, the mean spectra from *CaF*_2_ and *SS* showed much closer alignment (figure 3), indicating that despite *SS* producing stronger raw signals, both substrates captured similar underlying biochemical features. This normalisation step was essential to ensure that downstream machine learning models focused on spectral patterns rather than intensity artefacts.

**Figure 3.**
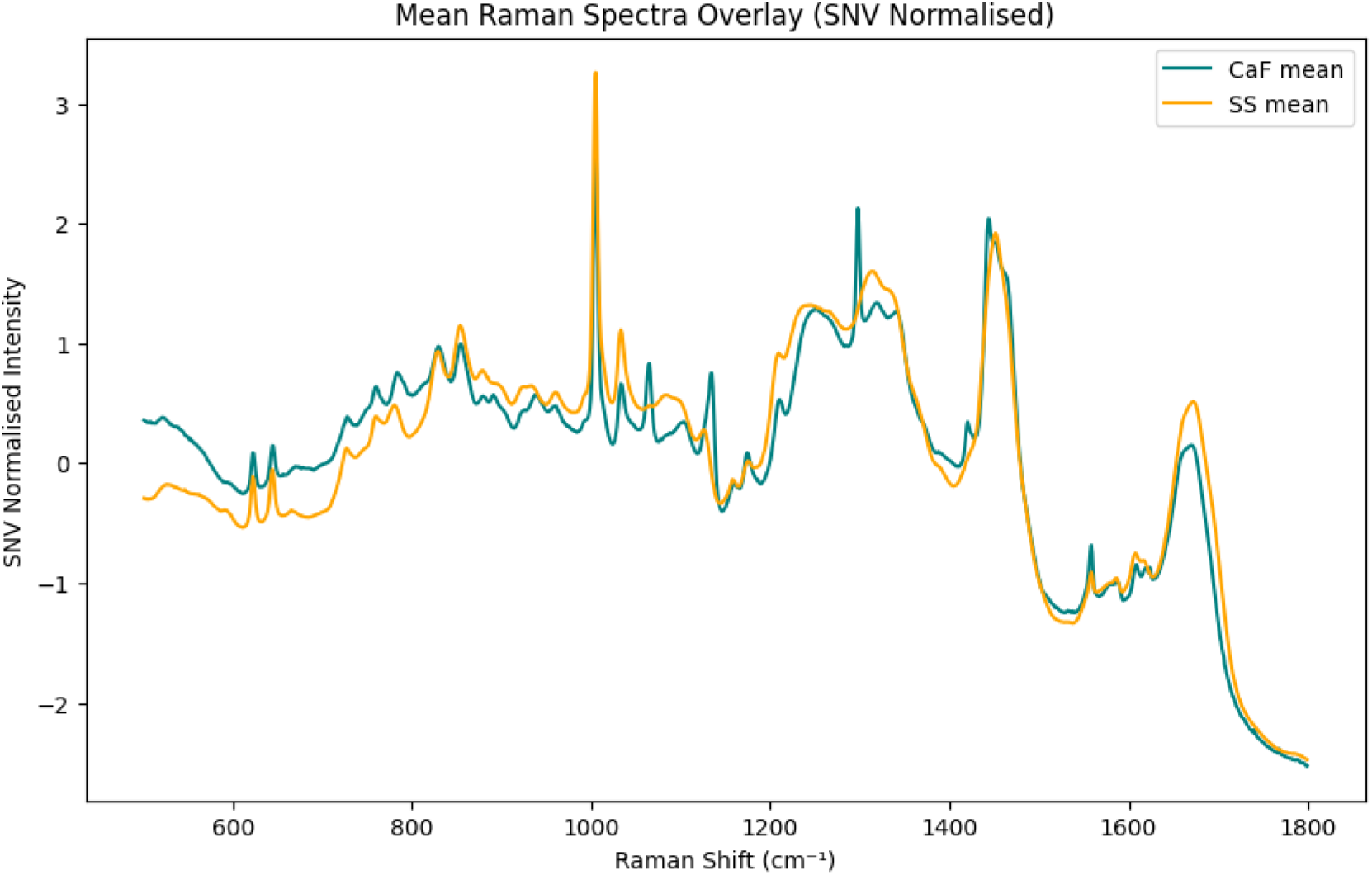
Mean normalised Raman spectra overlay from *CaF*_2_ and *SS* substrates.

### 3.2 Paraffin contamination and removal efficiency

Raman spectra acquired from *CaF*_2_ substrates revealed prominent paraffin contamination, with sharp peaks observed at 1062, 1132, 1170, 1295, 1417, 1439, and 1462 cm^*−*1^ (figure 4). These spectral features are characteristic of paraffin wax, a common embedding medium in FFPE samples, and are known to interfere with Raman-based tissue analysis^6^. The presence of these peaks indicates that standard dewaxing procedures were insufficient for complete paraffin removal on *CaF*_2_, leading to spectral interference.

**Figure 4.**
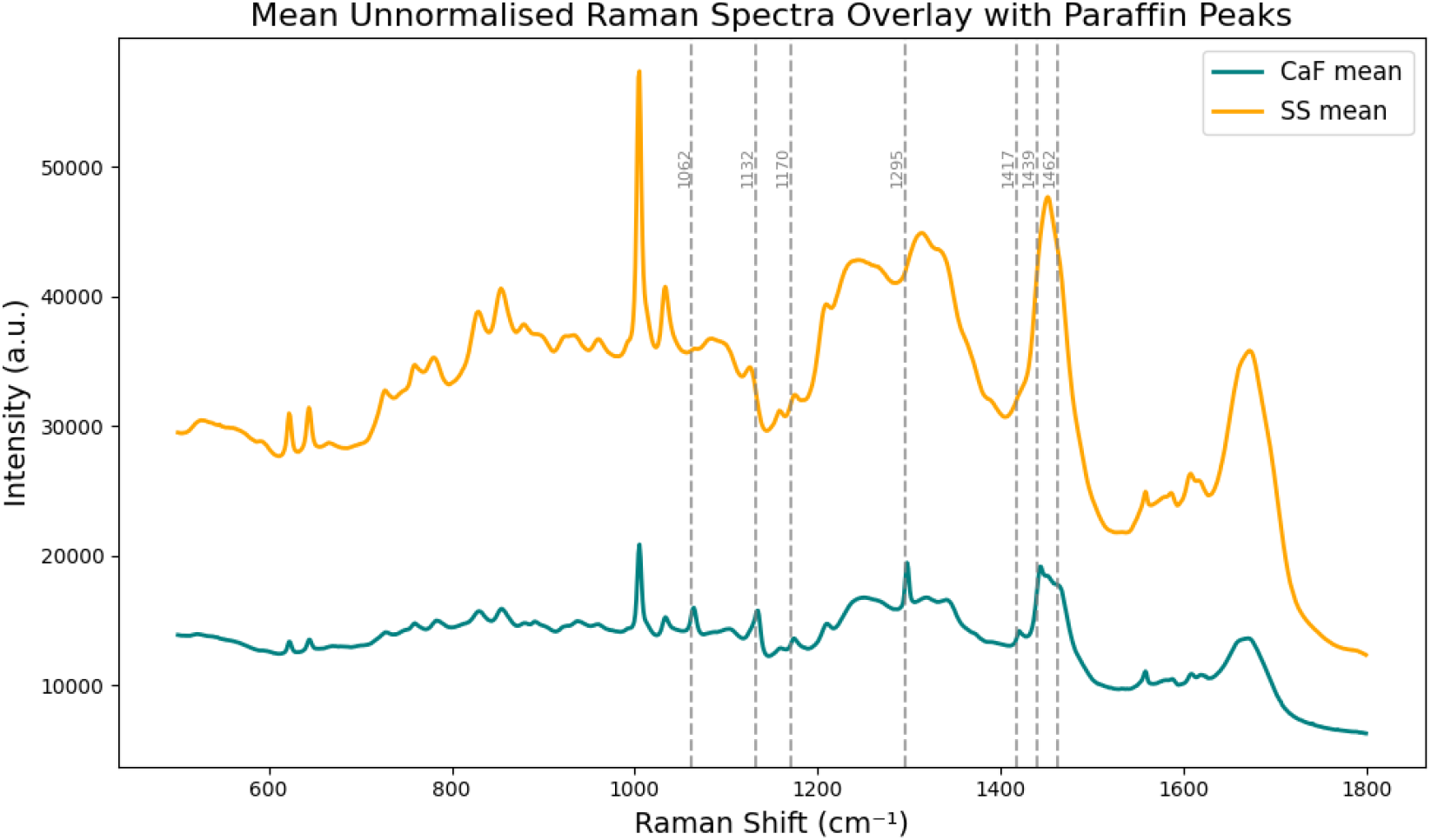
Mean unnormalised Raman spectra overlay from *CaF*_2_ and *SS* substrates, highlighting paraffin peaks.

In contrast, spectra collected from *SS* substrates showed minimal paraffin residues. This is likely due to *SS*’s favourable thermal and surface properties, which promote more effective paraffin removal ^6^. Previous work has demonstrated that *SS* supports thorough dewaxing across a range of FFPE tissues, even in archived samples.

To mitigate persistent paraffin contamination on *CaF*_2_, an extended dewaxing protocol was applied, involving an additional 48-hour xylene immersion. This reduced the average intensity of paraffin peaks by 10.8% (Table 1), leading to improved spectral clarity 5. However, some residual signals remain, potentially due to overlapping contributions from endogenous molecules such as lipids. Despite these improvements, the persistent presence of paraffin-associated peaks underscores the challenge of achieving complete dewaxing on *CaF*_2_.

**Table 1.** Reduction in paraffin peak intensity after extended dewaxing on *CaF*_2_.

| Peak (cm<sup>-1</sup>) | CaF Intensity | CaFII Intensity | Percentage Reduction (%) |
| --- | --- | --- | --- |
| 1062 | 0.759 | 0.648 | 14.62 |
| 1132 | 0.723 | 0.556 | 23.06 |
| 1295 | 1.624 | 1.614 | 0.60 |
| 1417 | 0.312 | 0.260 | 16.79 |
| 1439 | 1.774 | 1.663 | 6.30 |
| 1462 | 1.615 | 1.654 | 3.20 |

Since machine learning classifiers depend on biochemical spectral features, it is essential to minimise confounding signals from paraffin contamination. We hypothesised that residual paraffin peaks could impair classification performance by introducing non-biological variance. To investigate this, models were first trained on paraffin-contaminated spectra from the preliminary CaF dataset and later compared with models trained on spectra following extended dewaxing (CaFII). The following sections examine how paraffin interference influences the classification of ML-based tissue.

### 3.3 Machine learning classification - preliminary vs extended dewaxing

To assess the diagnostic utility of Raman spectra, machine learning classifiers were used to distinguish between EM and non-EM tissues (T1 CRC). Three models, PCA-LDA, SVM with RBF kernel, and GB, were evaluated. A nested cross-validation (CV) framework was employed, using `GroupKFold` in the outer loop (grouped by slide) and `StratifiedKFold` in the inner loop for hyperparameter tuning. This approach provides reliable performance estimates while preventing overfitting^8,9^.

As shown in Table 2, *SS* consistently achieved the highest classification accuracies across all models, while *CaF*_2_ initially underperformed due to paraffin interference. After implementing an extended dewaxing protocol, *CaF*_2_ accuracy markedly improved, especially for the SVM and GB classifiers, approaching the performance of *SS*.

**Table 2.**
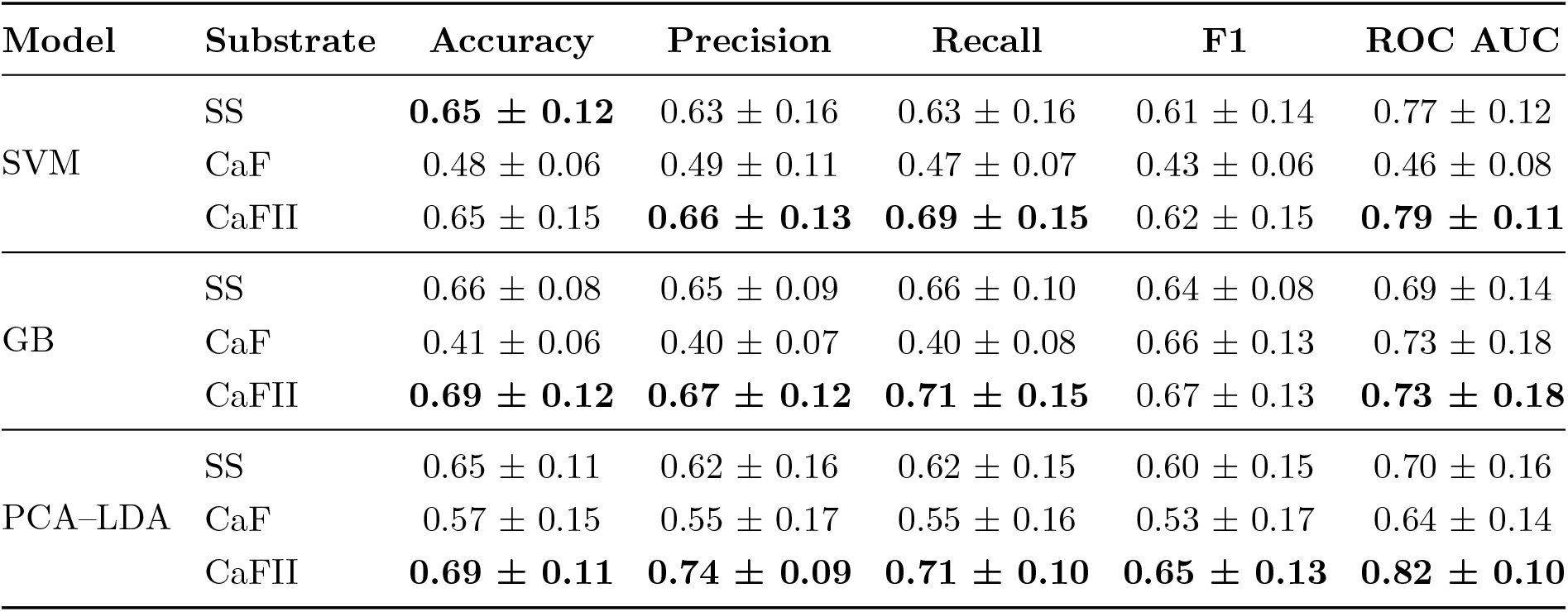
Performance metrics across substrates and ML models (mean ± SD, values rounded to two decimal places)

These results highlight two key findings: (1) *SS* offers inherently stronger diagnostic performance due to higher signal clarity and minimal paraffin retention; (2) extended dewaxing significantly enhances *CaF*_2_’s classification performance by reducing spectral noise from residual wax. Notably, *CaF*_2_ exhibited improved recall following extended dewaxing, which is clinically relevant for cancer diagnostics where sensitivity is important. This suggests that with adequate paraffin removal, *CaF*_2_ substrates can yield classification results comparable to *SS*, albeit with greatly increased increased sample preparation time (roughly 100-fold).

In summary, although stainless steel (*SS*) consistently demonstrates slightly superior diagnostic accuracy, efficiency, and spectral clarity (Table 2; Figure 5),the implementation of an extended dewaxing protocol clearly improves the performance of *CaF*_2_ by reducing paraffin interference and enhancing recall. These findings highlight that *CaF*_2_, when optimally prepared, achieves classification metrics largely comparable to *SS*, supporting the qualitative equivalance of this substrate. Notably, the increased sensitivity of *CaF*_2_ following extended dewaxing is advantageous in clinical scenarios where minimising false negatives is critical. Ultimately, substrate selection should be guided by the specific clinical or research context, considering factors such as cost, ease of sample preparation, and the diagnostic workflow.

**Figure 5.**
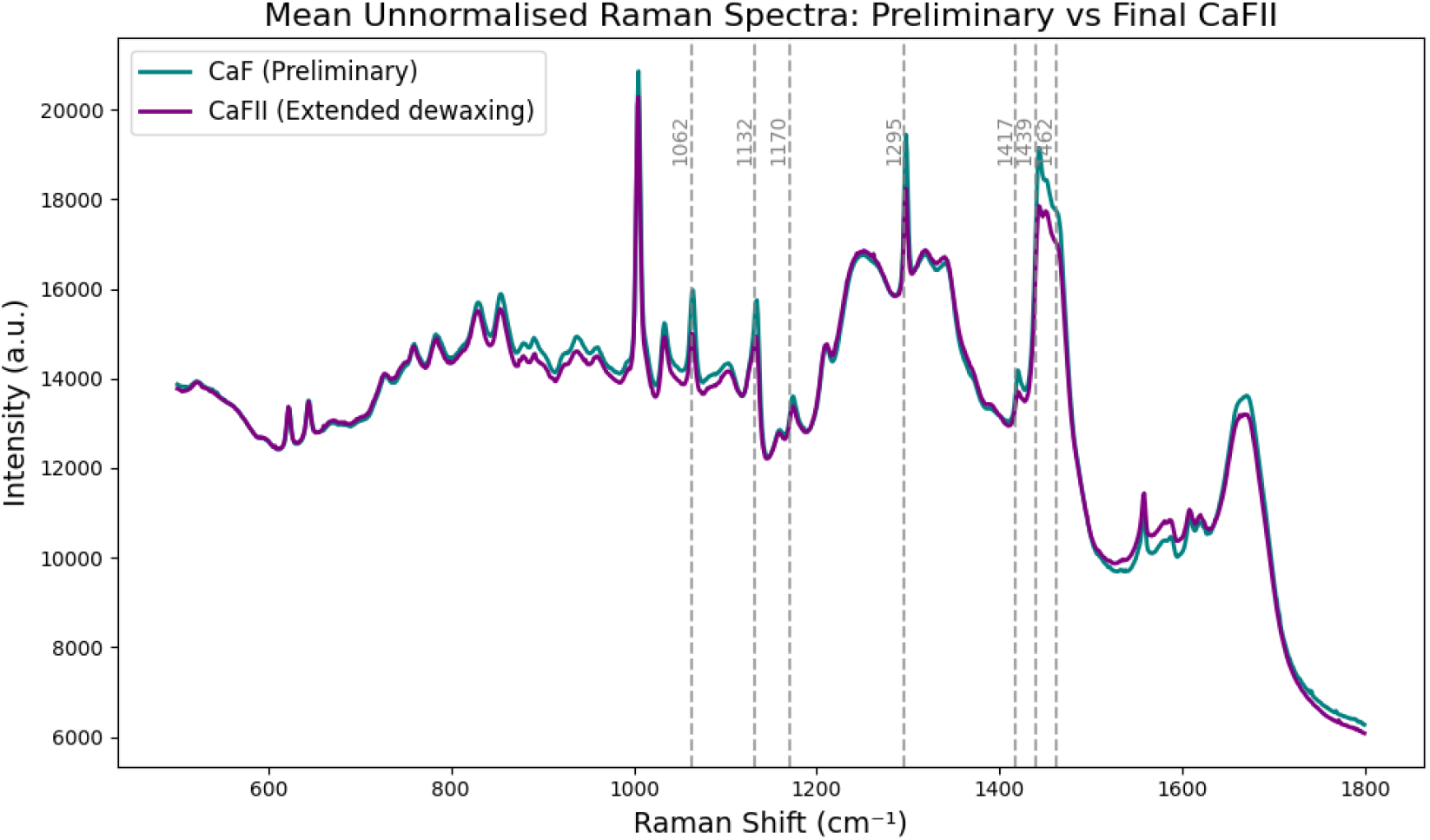
Mean unnormalised Raman spectra overlay from *CaF*_2_ (initial dewaxing) and CaFII (extended dewaxing) substrates, highlighting paraffin peaks.

## 4 Discussion

### 4.1 Ease of Preparation and Data Collection

Our findings demonstrate that mirrored stainless steel (*SS*) substrates offer significant practical advantages over calcium fluoride (*CaF*_2_) in Raman spectroscopy (RS). Previous studies have shown that *SS* facilitates faster spectral acquisition via a double-pass photon reflection mechanism, which improves light collection efficiency (including Raman-shifted photons) without compromising spatial resolution^5,6^. Additionally, *SS* shows minimal paraffin retention, simplifying and accelerating sample preparation by eliminating the need for prolonged dewaxing protocols.

By contrast, *CaF*_2_ substrates require higher laser power and extended acquisition times to achieve comparable signal intensity. Despite implementing an extended xylene immersion protocol (initially for 24 hours based on previous recommendations ^6^, followed by an additional 48 hours in this study, detailed in Supplementary 1), paraffin contamination persisted on *CaF*_2_ (Figure 5). This outcome was unexpected, as extended dewaxing was assumed to be sufficient for complete wax removal. Persistent paraffin required additional rounds of data collection, highlighting the challenge of obtaining clean spectral data from *CaF*_2_. In contrast, *SS* slides required only 40 minutes of dewaxing, which could perhaps even be shortened further.

### 4.2 Broader Implications for Clinical Workflows

#### 4.2.1 Cost-effectiveness and Scalability

*SS* substrates are markedly less expensive at approximately 4% of the cost of *CaF*_2_, making them far more suitable for scalable clinical use. This affordability supports high-throughput diagnostic workflows, particularly in modern pathology departments. Furthermore, *SS* substrates enhance Raman signal intensity by an average of 1.43 times (and theoretically up to fourfold), reducing acquisition time and increasing efficiency in both research and clinical settings^5^.

#### 4.2.2 Durability and Automation

Another key advantage of *SS* is its mechanical durability. Unlike brittle *CaF*_2_, which demands careful handling, *SS* is robust and better suited for storage, transport, and automation. This facilitates consistent and repeatable sample preparation, crucial for maintaining high-quality spectral output and enabling efficient automated RS workflows^10^.

#### 4.2.3 Signal Enhancement and Diagnostic Accuracy

The double-pass reflection mechanism in *SS* substrates leads to higher signal-to-noise ratio *CaF*_2_ ^5,6^. These enhanced spectra support more accurate tissue classification, particularly relevant in cancer diagnostics where subtle biochemical differences are critical. Notably, *SS* substrates demonstrate rapid wax melting under high laser power (Supplementary 2), evidencing their strong reflectivity. This reduces the need for high laser power and lowers the risk of tissue sample damage enabling confident use in further analysis with other techniques like immunohistochemistry. Consequently, *SS* substrates enable the development of robust machine learning (ML) models for automated diagnostics ^1^.

#### 4.2.4 Integration with AI-driven Diagnostics

High-quality, reproducible data from *SS* should enhance compatibility with AI-driven diagnostic models. Advanced ML and deep learning (DL) algorithms are increasingly applied to Raman spectra for cancer detection^1,9^. The superior spectral clarity of *SS* improves model performance by reducing noise and variability, ultimately supporting earlier and more accurate diagnoses. This aligns with ongoing shifts towards digital pathology, where cost-effective and scalable platforms are essential^11^.

### 4.3 Limitations and Future Directions

Our acquisition parameters were optimised for *CaF*_2_, potentially underutilising the signal-enhancing capabilities of *SS*. Future studies should design protocols specifically optimised for *SS*. We also note that *CaF*_2_ performance which improved with extended dewaxing may decline rapidly with suboptimal preparation.

The dataset included only 35 colorectal tissue samples, which may limit at this early stage the generalisability of findings to other cancers. A larger, more diverse dataset is necessary to validate these conclusions across tissue types. Additionally, only traditional ML models were used, as the dataset size did not support DL approaches. With larger datasets, DL could uncover more complex spectral patterns and enhance classification accuracy^1,8^.

## 5 Conclusion

This study demonstrates that mirrored stainless steel (*SS*) substrates offer several advantages over calcium fluoride (*CaF*_2_) for Raman-based digital pathology in colorectal cancer. *SS* substrates provide enhanced signal intensity, superior reproducibility, and significantly lower cost, while requiring simpler and faster sample preparation.

Although extended dewaxing protocols improve *CaF*_2_ performance by reducing paraffin contamination and increasing sensitivity, *SS* still outperforms in diagnostic accuracy, efficiency, and practical utility. Indeed, it is conceivable that *CaF*_2_ might not have produced any useful classification if the comparision between substrates had been made using conditions optimised for SS rather than *CaF*_2_. These findings underscore the importance of substrate selection and optimised preparation protocols in Raman-based diagnostics and support the broader integration of *SS* into AI-driven, scalable clinical workflows.

## Author contributions

Conception: NB, GT. Development of methodology: NB, GT. Data acquisition: CN. Data analysis: CN, NB. Funding acquisition: GT Manuscript writing. NB, CN

## Data availability

Data is available via Zenedo: https://zenodo.org/records/17493210.

## Ethics Statement

Consent for using human tissues was obtained via UCL/UCLH Biobank for studying Health and Disease (REC 25/YH/0120).

## Acknowledgements

Thanks to Professor Manuel Rodriguez-Justo at Department of Research Pathology, Cancer Institute, UCL for clinical support and Riana Gaifulina for technical support.

This work benifitted from funding from an EPSRC Ph.D. Studentship (EP/R513143/1)and seed funding from EPSRC CLIRPath-AI seedfunding (EP/W00058X/1).

## Supplementary Data

### Supplementary 1. Paraffin Removal Protocol

All tissue slides underwent a 24-hour incubation at 37°C prior to dewaxing. Dewaxing was performed via four successive xylene baths (VWR International Ltd., UK) with gentle agitation, followed by graded ethanol rehydration (absolute ethanol, VWR International Ltd., UK). Ethanol rehydration consisted of two sequential immersions in 100%, 90%, 70%, and 50% ethanol for five minutes each, followed by a final water rinse (Table S1).

**Table S1.** Dewaxing protocols for tissue sections on different substrates.

| Slide Type | Dewaxing Protocol |
| --- | --- |
| All Slides | 4x xylene baths with agitation; 2x immersions each in 100%, 90%, 70%, 50% ethanol for 5 min; final water rinse. |
| Stainless Steel (SS) | Xylene 1–4: 10 min each; 100% EtOH (2x): 5 min; 90% EtOH (2x): 5 min; 70% EtOH (2x): 5 min; 50% EtOH (2x): 5 min; Water: 10 min. |
| $CaF_2$ (CaF) | Xylene 1–3: 10 min; Xylene 4: overnight (24 hrs); same ethanol steps as above. |
| Extended Dewaxed $CaF_2$ (CaFII) | As CaF above, with additional: Xylene 1–3: 10 min; Xylene 4: extended to 48 hrs; ethanol steps as above. |

### Supplementary 2. Laser-induced effects on substrates

**Figure S1.**
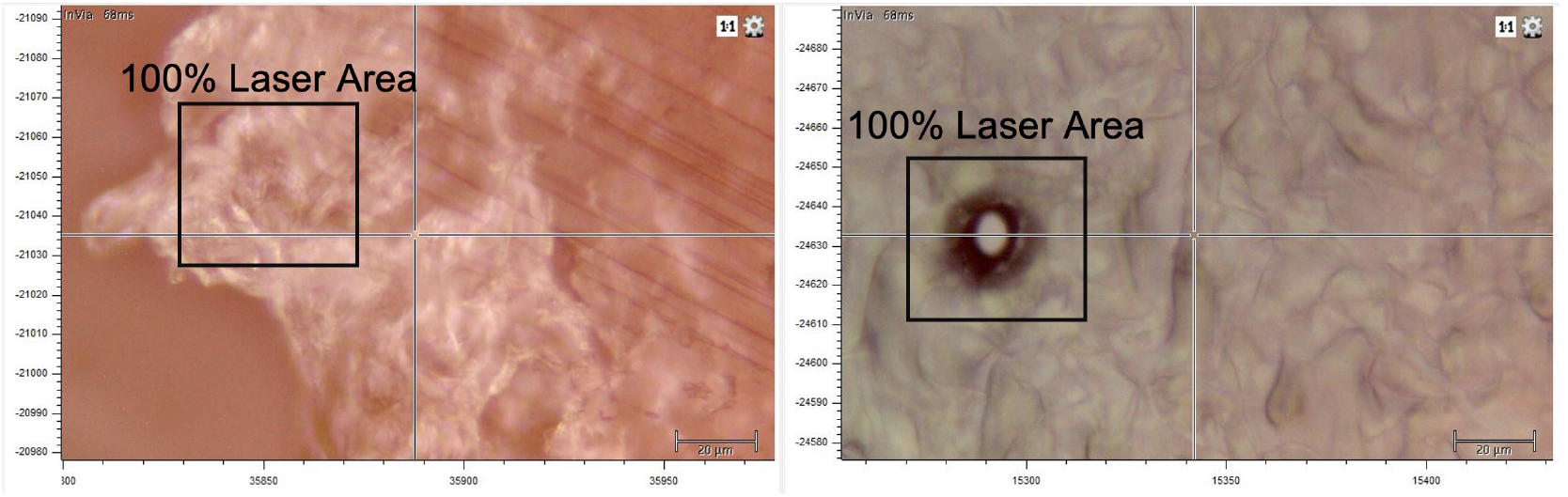
Representative optical micrographs showing pure paraffin and laser-induced effects on *CaF*_2_ (left) and SS (right) substrates under 10% and 100% laser exposure. The SS substrate (right) exhibits a localised “burnt” area where the laser was applied at 100% power.

### Supplementary 3. Optimised stainless steel data acquisition

To assess whether stainless steel (*SS*) substrates retain their superior performance under accelerated acquisition settings, we conducted a rapid optimisation protocol. Unlike the standard protocol using 10 accumulations, spectra were acquired with only 1 accumulation and minimal adjustment, simulating a fast, high-throughput workflow. This allowed us to evaluate whether *SS* still delivers usable spectral quality when experimental time is constrained.

**Figure S2.**
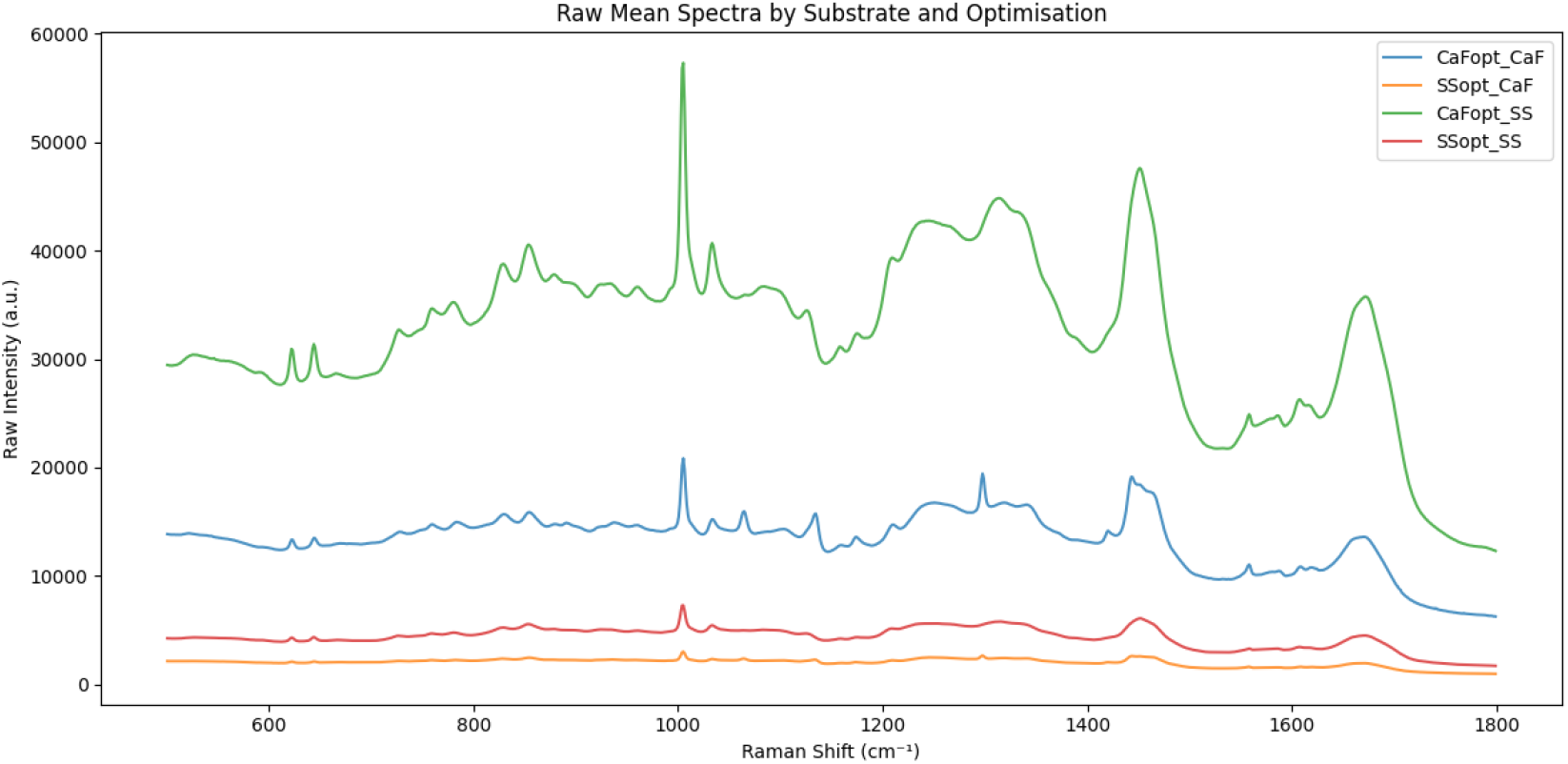
Comparison of SNV-normalised Raman spectra between stainless steel (SS) and extended-dewaxed *CaF*_2_ (CaF_2-opt_). SS spectra show enhanced clarity and stronger biochemical band resolution.

